# Regularized Deep Neural Networks for Combining Heterogeneous Features of Peptides in Data Independent Acquisition Mass Spectrometry

**DOI:** 10.1101/2025.06.09.658564

**Authors:** Namgil Lee, Hojin Yoo, Dohyun Han, Heejung Yang

## Abstract

Data-independent acquisition (DIA) has gained much attention in mass spectrometry (MS)-based proteomics for its improved reproducibility and unbiased data acquisition. In DIA-MS, the spectral library is crucial in peptide identification. However, this method is limited to peptides previously identified via data-dependent acquisition (DDA) MS experiments. This study proposes a deep learning approach for generating spectral libraries, even for previously unseen peptides. While most deep learning-based methods rely on one-hot encoding representation for peptides, the proposed method incorporates physicochemical features, including atomic composition, hydrophobicity, flexibility, fractional surface probability, and aromaticity. We introduce sparsity regularized neural network layers to facilitate the selection and combination of important high-dimensional physicochemical features and improve prediction performance. Furthermore, we suggest a transfer learning strategy for training the proposed deep neural networks having multiple heterogeneous input channels. Numerical experiments using benchmark DDA-MS data demonstrated that the proposed deep learning model out-performed existing benchmark models, such as Prosit and DeepDIA, particularly in predicting retention times. And it was demonstrated that the proposed models with sparsity regularization identified more peptides from HeLa cell DIA data compared to the other deep learning models.

## Introduction

Mass spectrometry (MS) has become a standard tool in proteomic research for the proteomic analyses of biological systems^1,2^. In liquid chromatography-tandem MS (LC-MS/MS), a protease digests proteins into peptides, which are further fragmented into fragment ions. The relative abundance of the fragment ions is then measured in the form of mass spectra. One of the most widely used data acquisition strategies for tandem MS is data-dependent acquisition (DDA) strategy: only a subset of precursor ions selected from the MS survey scan is processed in subsequent MS scans to obtain tandem MS spectra. In contrast, a data-independent acquisition (DIA) strategy has been introduced more recently as an accurate and reproducible method for detecting large fractions of peptides^3^. The DIA strategy collects tandem MS spectra from a series of MS scans covering broad precursor ion ranges, guaranteeing unbiased and consistent data collection.

For peptide identification and quantification, a DIA strategy requires fragment ion spectral libraries generated for target peptides. The fragment ion intensities obtained from current experimental data are correlated with a spectral library for peptide identification ^4^. Hence, the quality of spectral libraries is crucial for accurate peptide identification and quantification in DIA MS. Traditional methods for generating spectral libraries depend on fragment ion spectra obtained from the DDA data. Because of this dependence, the performance of the generated spectral libraries can be hindered by a mismatch in the instrument type, limited coverage of DDA samples, or high false discovery rates^5^.

To overcome the limitations of traditional proteomics approaches based on MS, researchers have developed deep learning methods for various aspects of the proteomics workflow. Deep learning methods for predicting fragment ion intensities and retention times have been developed and applied to in silico spectral library generation^6^. An advantage of deep learning, and more generally machine learning, is that the inferred models can be generalized to new peptides. For example, peptides originating from another organism can be predicted^7^. However, traditional machine learning methods use a limited number of known features for peptide representation. For instance, each peptide is represented by a 20-dimensional vector consisting of 20 amino acid residues^8^. Because unknown factors and structural information on peptides can affect the measured spectra, using only known features can cause high prediction errors.

In contrast, modern deep learning methods, such as convolutional neural networks (CNN) and recurrent neural networks (RNN)^9^, have been adopted to build models that can detect peptide features automatically. Because of the sequence structure of peptides, most popular deep learning methods are based on neural networks originally developed for natural language processing, such as the RNN^10^, long short-term memory (LSTM)^11^, and gated recurrent unit (GRU)^12,13^. LSTM and GRU are suitable for modeling short- and long-range interactions between the amino acids in peptides. Many deep learning methods for fragment ion intensity (MS2) prediction have been developed based on bidirectional LSTM (or GRU) and onehot encoding representations for peptides^14–19^. Using a one-hot encoding representation, scientists have encoded each amino acid as a 20-dimensional vector of zero-one values. Deep learning models automatically extract latent features via gradient-based optimization of model parameters.

However, the one-hot encoding representation of peptides has potential drawbacks. First, many amino acid modifications exist; thus one-hot encoding representation cannot be generalized to potentially unseen modified peptides. Second, it may not produce chemically or physically interpretable latent features in deep learning models. Third, the performance of prediction models depends heavily on the data quality and computational power, which ignores prior knowledge of the chemical properties of peptides. Alternatively, a CNN-based deep learning method, MS2CNN, converts peptide sequences into feature vectors of peptide properties, such as amino acid composition, secondary structure, and physicochemical features^20^. In addition, to improve the generalization ability of deep learning models to previously unseen modified peptides, Bouwmeester et al. ^21^ proposed using peptide features based on atomic composition.

Although all the studies mentioned above have proposed featurization methods for peptides and amino acids, combining various features to achieve high performance and physiological interpretation is worthwhile. This paper proposes a regularization method to combine one-hot encoding features with high-dimensional physicochemical features of peptides. The advantages of the proposed method are: first, using one-hot encoding features can enable deep learning models to extract latent features from data automatically. Second, the generalization performance of the deep learning model can be dramatically improved for modified peptides not included in the training data using peptide features based on atomic composition. Third, sparsity regularization prevents overfitting and selects useful features for MS spectral prediction, which improves interpretability of model performances. Fourth, high-dimensional heterogeneous features can be combined efficiently by a transfer learning strategy for training deep neural networks having multiple input channels. Figure 1 illustrates the proposed in silico spectral library generation method based on sparsity regularized deep neural networks.

**Figure 1.**
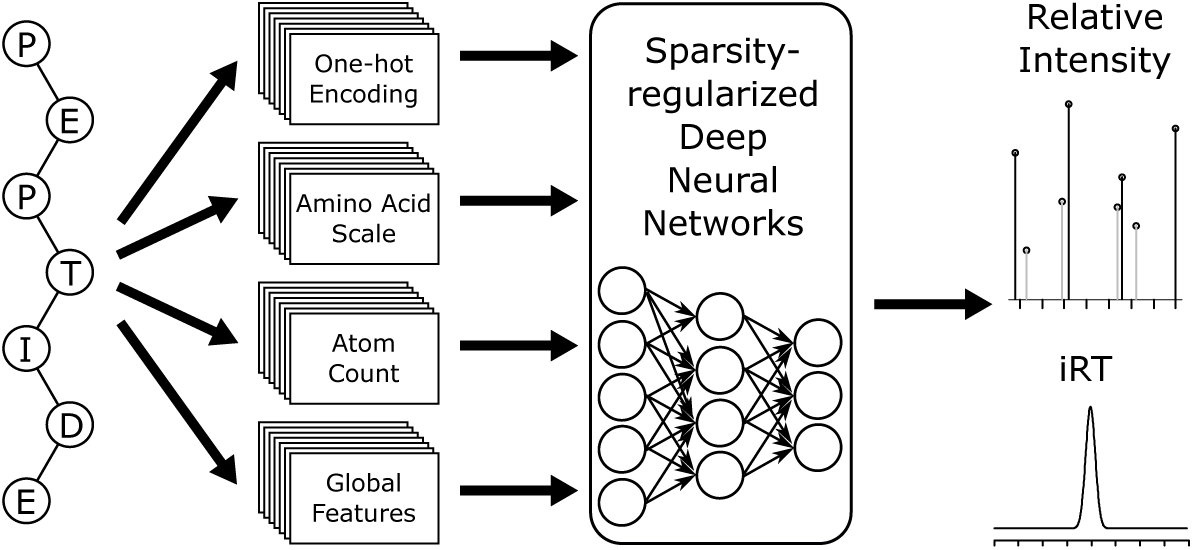
: Introduction to the proposed sparsity regularized deep neural networks for generating in silico spectral libraries.

## Methods

Two deep learning models are required separately to generate a deep learning based spectral library: one for predicting fragment ion intensities and the other for predicting indexed retention time (iRT). In this work, both models are constructed consisting of featurization layers, feature extraction layers, a concatenation layer, and a fully connected layer, as depicted in Figure 2. A more detailed description is provided below:

**Figure 2:**
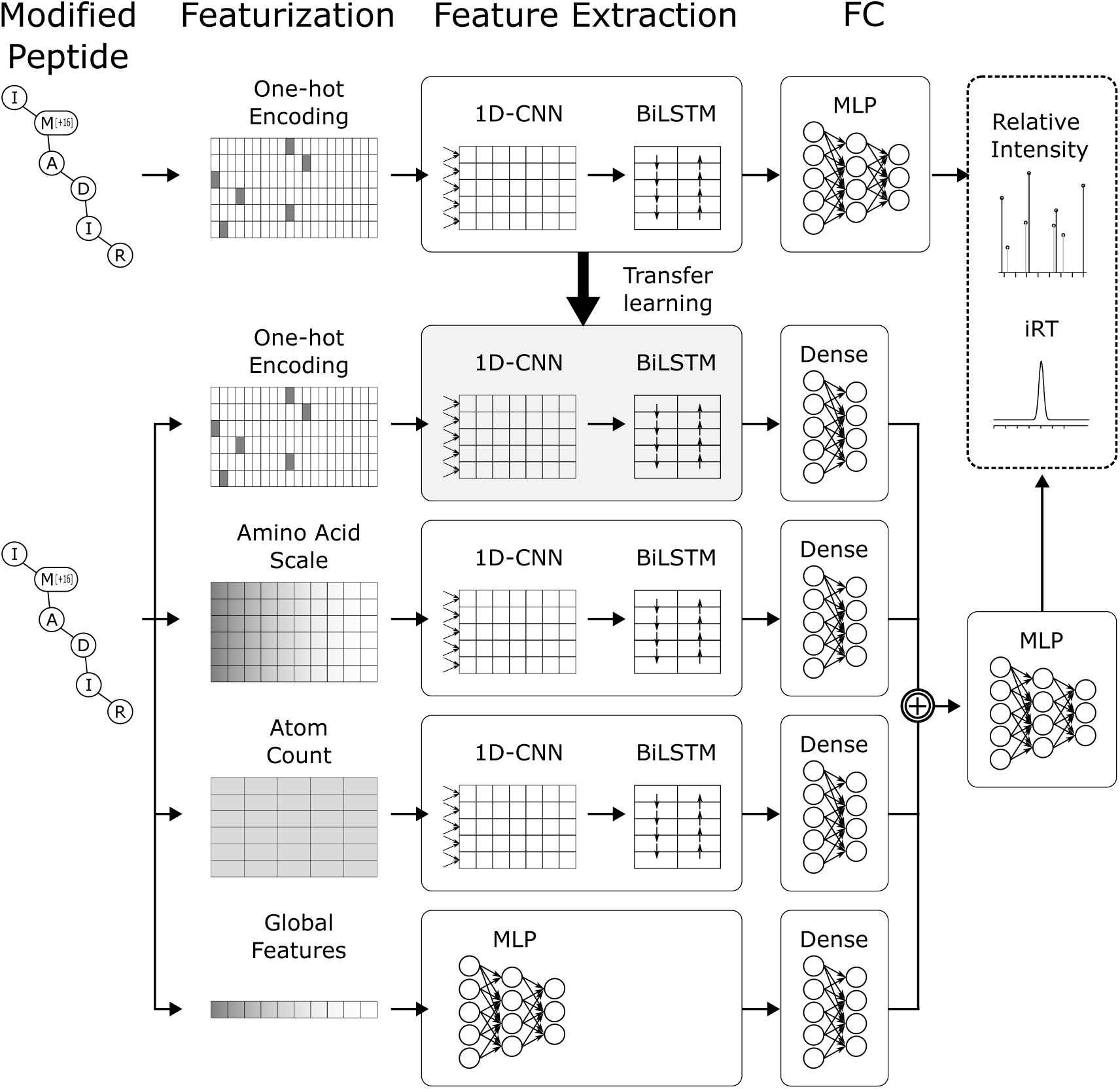
Architecture of the proposed deep learning models

### Heterogeneous Features of Peptides

The proposed deep learning models use modified peptide sequence as input. The featurization layers consist of multiple parallel converters for extracting heterogeneous numerical features from the peptides. The features can be grouped into four categories: each is forwarded to a proper deep neural network pipeline in the feature extraction layers to extract higher-level features at deeper layers.

#### One-hot Encoding Features

One-hot encoding is a featurization technique motivated by natural language processing. A sentence is represented as a sequence of words; likewise, a peptide can be represented as a sequence of amino acids. Considering the dictionary of the 20 standard amino acids, onehot encoding of a peptide converts each amino acid in the peptide into a zero-one-valued vector of length 20, where zeros and ones indicate the existence of a specific amino acid. In addition, because peptide sequences vary in length, the length difference is adjusted to fixed length through zero padding so that an RNN-based deep learning model can be applied in any subsequent layer. Consequently, the one-hot encoding of a peptide is represented as a zero-one matrix of size *L* × *A*, where *L* is the zero-padded peptide length and *A* is the number of amino acids in the dictionary.

Although one-hot encoding has been widely adopted in deep learning approaches, it has limitations in generalizing to a wide variety of peptide modifications. Combining physicochemical features with one-hot encoding features can improve the generalization performance of deep learning models.

#### Amino Acid Scale Features

The amino acid scale features are the physicochemical properties of each amino acid, such as hydrophobicity^22^. While there are many predefined scales introduced in the literature, we chose five representative scale features to be incorporated in the numerical experiments: (1) the Kyte & Doolittle index of hydrophobicity^23^, (2) Wilson hydrophobic constants^24^, (3) flexibility parameters^25^, (4) fractional surface probability^26^, and (5) interior-to-surface transfer energy scale^27^. A peptide’s amino acid scale features are represented as a matrix of size *L* × *S*, where *L* is the zero-padded peptide length and *S* is the number of chosen scale features.

#### Atom Count Features

Atom count features represent the atomic composition of each amino acid. Basic atom count features were used to calculate the numbers of five atoms (C, H, N, O, and S) in each of the 20 standard amino acid types. The numbers of added and deleted atoms were calculated separately and integrated into the total number of atoms for modified amino acids. Consequently, the atom count features of a modified peptide are a matrix of size *L* × 5. Computing the atom count features is straightforward for any peptide; thus, any model using atom count features can be generalized to a modified amino acid with a known atomic composition.

#### Global Features

Global features encode the physicochemical properties of a peptide^22^. We retrieved 16 global features of a peptide using the SeqUtils module in the Biopython package^28^: length, molecular weight, aromaticity, charge at pH 5.0, charge at pH 7.5, minimum flexibility, median flexibility, maximum flexibility, grand average of hydropathy ^23^, instability index^29^, isoelectric point^30^, molar extinction coefficient assuming cysteine (reduced), molar extinction coefficient assuming cysteine residues (Cys-Cys bond), the fraction of amino acids in the helix, the fraction of amino acids in the turn, and the fraction of amino acids in the sheet. The global features of a peptide are represented with a vector of length 16.

### Regularized Neural Networks for Heterogeneous Data Fusion

Multiple heterogeneous features obtained in the featurization layers are forwarded to the deep neural network layers for higher-level feature extraction. Specifically, pipelines of deep neural network layers exist for processing each type of heterogeneous feature separately. The pipelines of deep neural network layers can be categorized into two types according to the input feature size. One type is for matrix-shaped features and the other is for vector-shaped features.

Matrix-shaped features can encode sequence information of the peptide by preserving the positions of every amino acid in the sequence. In this case, the deep neural network layers consist of 1D CNN layers and bidirectional long short-term memory (BiLSTM) layers motivated by DeepDIA^19^. The kernel sizes of the 1D CNNs were set to two such that the CNN kernels could extract local feature patterns of the two neighboring amino acids from the whole sequence structure^9^.

Conversely, vector-shaped features can encode the global properties of peptides rather than sequence-structure information. The deep neural network layers for vector-shaped features consist of fully connected layers with a rectified linear unit (ReLU) as the activation function. With dropout layers placed between the fully connected layers, the resulting deep neural network layers form a multilayered perceptron representing a nonlinear transformation of the input features.

Notably, the input features are heterogeneous and high-dimensional; thus, redundant features may have been included. L1-norm regularization is known to prevent overfitting and to yield sparse weights^31^. Hence, for each pipeline of deep neural network layers described above, the weights of the first layer were regularized by the L1-norm, except the pipeline for one-hot encoding features. In other words, let *W* and *b* denote the kernel weights and bias of the first CNN layer or the first fully connected layer in a deep neural network pipeline. Then, during the training of the model, an optimizer minimizes the L1-norm regularized loss function, which can be expressed as

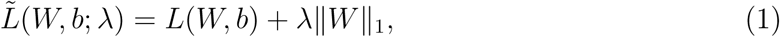

where *L*(*W, b*) is a loss function, *λ* ≥ 0 is a regularization parameter, and ∥*W*∥_1_ = ∑_*i*_|*w_i_*| is the L1-norm of weight *W* = (*w_i_*). The regularization parameter *λ* can be set differently for each deep neural network pipeline such that each type of feature can be weighted adaptively.

### Transfer Learning

Because the proposed model consists of multiple deep neural network pipelines corresponding to heterogeneous input channels, gradient-based learning algorithms suffer from local optima and poor weight initialization. To mitigate such problems, we suggest a transfer learning strategy, which is depicted in Figure 2. First, a deep neural network consisting of a featurization layer, a feature extraction layer, and a fully connected layer is constructed, where the featurization layer extracts only one-hot encoding features from peptides. Second, hyperparameters of the deep neural network are optimized and the network weights are trained. Third, the weights of the feature extraction layer are fixed and transferred to an another deep learning model, called Fusion, consisting of multiple deep neural network pipelines. Finally, hyperparameters of the Fusion model are optimized and the network weights are trained, except the weights of the transferred feature extraction layer corresponding to the one-hot encoding features.

### Hyperparameter Optimization

We compared several deep learning models to generate in silico spectral libraries. The deep learning models used in this study can be categorized into (1) MS2 prediction models and (2) iRT prediction models. Note that the MS2 prediction models were trained separately on the precursor charges of +2 and +3 values. A charge 2 model and a charge 3 model may have different sets of hyperparameters even if they consist of the same types of deep learning layers. The names and brief description of the models are as follows.

(A) Prosit: Pretrained Prosit models^15,32^.

- MS2: A Prosit model released in 2020 with higher energy C trap dissociation (HCD) fragmentation^32^.
- iRT: A Prosit model released in 2019^15^.
(B) Baseline: Deep learning models provided by DeepDIA^19^.

- MS2: The input features consisted of only one-hot encoding features, and the model consisted of a single 1D-CNN layer, a single BiLSTM layer, and a single fully connected layer.
- iRT: The model used one-hot encoding features as input and consisted of a 1DCNN layer, a max-pooling layer, a BiLSTM layer, and three fully connected layers.
(C) CRNN: Extension to the DeepDIA.

- MS2: The input features consisted of only one-hot encoding features, and the model consisted of multiple 1D-CNN layers, multiple BiLSTM layers, and multiple fully connected layers.
- iRT: Extension to the DeepDIA. The model consisted of multiple 1D-CNN layers, a max pooling layer, multiple BiLSTM layers, and multiple fully connected layers.
(D) Fusion: The proposed deep learning model using multiple deep neural network pipelines corresponding to heterogeneous input channels. As explained in Section Transfer Learning, the weights of the feature extraction layer of the CRNN model were transferred to the pipeline for the one-hot encoding features. The first layer of each of the other pipelines was L1-norm regularized.
(E) Fusion w.o. L1: The Fusion model without L1-norm regularization.
(F) Fusion w.o. scale: The Fusion model without amino acid scale features.

For hyperparameter optimization, the entire training set was split into training and validation sets with split ratios of 67% and 33%, respectively. The best set of hyperparameters was selected at the minimum of the validation set losses: the mean squared error (MSE) for the MS2 prediction models and the mean absolute error (MAE) for the iRT prediction models. See Supporting Information for detailed model architectures.

Hyperparameter optimization and numerical experiments were performed on a workstation with an Intel CPU (3.00 GHz) and an NVIDIA GPU (GeForce RTX 3060 Ti) running Ubuntu 18.04. All deep learning models were implemented in Python 3.9 and Tensorflow 2.4. Hyperparameter optimization was performed using the Comet machine learning platform (https://www.comet.com/) with the Python package comet-ml 3.1.

## Results

### Data Preparation

The LC-MS/MS DDA data from four experiments were used to generate experimental spectral libraries. Each of these libraries was subsequently used to train deep learning models, with the HeLa data specifically utilized for hyperparameter optimization. A summary of the generated spectral libraries are provided in Table 1.

**Table 1:** Summary of the experimental spectral libraries used for training the deep learning models.

| Dataset | Protein Groups | Modified Peptides | Precursors | Fragments | Dataset Identifier |
| --- | --- | --- | --- | --- | --- |
| HeLa | 6,088 | 69,515 | 96,354 | 1,187,745 | PXD005573 |
| Mouse cerebellum | 10,062 | 171,831 | 258,835 | 6,048,675 | PXD014108 |
| Pombe (chymotrypsin) | 1,191 | 5,379 | 6,289 | 150,011 | PXD011459 |
| Pombe (trypsin-chymotrypsin) | 1,783 | 17,024 | 19,623 | 523,415 | PXD011459 |

First, the DDA data from HeLa cells were acquired on a Q Exactive HF mass spectrometer (Thermo Fisher Scientific, Waltham, MA, USA), available at ProteomeXchange (proteomecentral.proteomexchange.org/), with the dataset identifier PXD005573^33^. The 17 raw file names begin with C_D160304_S251-Hela-2ug-2h_MSG, C_D160331_S209-HPRP-HeLa, and C_D160401_S209-HPRP-HeLa. The DDA data were analyzed using Spectronaut 15^34^ with default settings, except that Carbamidomethyl (C) was selected as the fixed modification, and no other modifications were applied. The DDA data were searched against the UniProt Homo sapiens FASTA database (https://www.uniprot.org/, accessed on 2021-09-13, containing 20,588 entries).

In addition, LC-MS/MS DIA data for HeLa cells were obtained from the same repository of ProteomeXchange with the dataset identifier PXD005573. The DIA data were technical replicates acquired using the same experimental settings, which might be useful for validating the reproducibility of peptide identification. The three raw file names begin with Fig1_MP-DIA-120min120kMS1-22W30k-8dppp.

Second, the experimental spectral library generated from Mouse cerebellum experiments was obtained directly from ProteomeXchange with the dataset identifier PXD014108. The original raw DDA data files were acquired on a Q Exactive HF mass spectrometer from 15 fractionated runs and are available at ProteomeXchange with the dataset identifier PXD005573. The raw files were searched using SpectroMine (version 1.0.21621) with the default settings, against the SwissProt Mus musculus FASTA database (accessed on 2019-02, containing 17,006 entries). The generated spectral library is available in the zipped file mouse_cerebellum_DDA.csv.zip. All cysteine residues in the peptide sequence were considered modified by carbamidomethylation. See Yang et al. ^19^ for more details on the database searching of the “Mouse2” DDA data.

Lastly, the DDA data files from S. pombe cells were acquired on an Orbitrap Elite mass spectrometer (Thermo Fisher Scientific, Waltham, MA, USA) and are available at ProteomeXchange with the dataset identifier PXD011459. The S. pombe cell lysates were processed either with chymotrypsin alone, or with trypsin followed by chymotrypsin. The acquired DDA data files were searched against the PombeBase FASTA database (https://www.pombase.org/, accessed on 2024-01, containing 47,432 entries) using MaxQuant (version 2.4.10.0) ^35^. The MaxQuant parameters were set as follows: Carbamidomethyl (C) as a fixed modification, Oxidation (M) as a variable modification, MS accuracy of 4.5 ppm, MS/MS tolerance of 0.5 Da, and a maximum number of missed cleavages set to 6. See Dau et al. ^36^ for more details.

### Numerical Evaluation

The deep learning models were trained and tested using the experimental spectral libraries described in the previous section. Precursors with charge states greater than 3 or less than 2 were removed for training the deep learning models because their number was much smaller than that of the other precursors, which could lead to highly biased training and validation sets. To maintain the data split ratio identical to that of the hyperparameter optimization step, we split the entire training set into a training set and a validation set with a 2:1 ratio. The models were then numerically compared based on the validation set.

#### HeLa Data

Table 2 summarizes the performance of deep learning models in terms of the MSE, MAE, and mean absolute percentage error (MAPE). The Fusion w.o. L1 and CRNN models yielded the lowest MSE and MAE values among the MS2(+2) prediction models, whereas the Baseline model showed overall low performance because its architecture had not been optimized. The CRNN model yielded the lowest MSE and MAE values among the MS2(+3) prediction models, which implies that a relatively simpler model was sufficient for predicting the MS2 of precursors with charge state +3 than with charge state +2. On the other hand, the Baseline and Fusion models yielded the lowest MAPE values, which is due to their relatively simple architectures and L1-norm regularization that enhanced robustness to outliers. The Fusion w.o. L1 model achieved the lowest MSE and MAE values among the iRT prediction models, demonstrating that physicochemical features offer a significant advantage for iRT prediction. The pretrained Prosit models yielded the worst performance because they had been trained on different datasets.

**Table 2:** Comparison of in silico spectral library generation models trained and validated on HeLa data. The MSE, MAE, and MAPE are reported with standard errors listed below each one. The bold font indicates the smallest number among the models.

| Model type |  |  | Prosit | Baseline | CRNN | Fusion | Fusion w.o. L1 | Fusion w.o. scale |
| --- | --- | --- | --- | --- | --- | --- | --- | --- |
| MS2<br>(+2) | MSE | $\times 10^{-4}$ | 17.40 | 5.90 | 3.23 | 4.74 | **3.17** | 3.41 |
| | | $\times 10^{-6}$ | (8.50) | (3.20) | (2.44) | (3.65) | (2.44) | (2.53) |
| | MAE | $\times 10^{-3}$ | 4.77 | 2.66 | **1.88** | 2.09 | 1.90 | 1.93 |
| | | $\times 10^{-5}$ | (1.25) | (0.73) | (0.54) | (0.65) | (0.53) | (0.55) |
| | MAPE | $\times 10^{-2}$ | 2.87 | 1.27 | 1.28 | **1.14** | 2.26 | 1.64 |
| | | $\times 10^{-5}$ | (4.93) | (3.20) | (3.13) | (2.94) | (4.27) | (3.60) |
| MS2<br>(+3) | MSE | $\times 10^{-4}$ | 24.61 | 9.40 | **4.92** | 5.46 | 6.43 | 5.74 |
| | | $\times 10^{-6}$ | (13.92) | (6.05) | (4.06) | (4.78) | (5.49) | (4.95) |
| | MAE | $\times 10^{-3}$ | 6.97 | 3.82 | **2.72** | 2.79 | 2.95 | 2.86 |
| | | $\times 10^{-5}$ | (2.16) | (1.32) | (0.95) | (1.01) | (1.09) | (1.03) |
| | MAPE | $\times 10^{-2}$ | 4.48 | **1.72** | 1.74 | 2.15 | 2.46 | 2.68 |
| | | $\times 10^{-5}$ | (9.16) | (5.43) | (5.26) | (5.93) | (6.42) | (6.70) |
| iRT | MSE | $\times 10^{-4}$ | 322.78 | 6.03 | 5.92 | 6.06 | **5.60** | 5.94 |
| | | $\times 10^{-5}$ | (30.87) | (4.35) | (4.29) | (4.31) | (4.29) | (4.25) |
| | MAE | $\times 10^{-2}$ | 16.83 | 1.13 | 1.20 | 1.19 | **1.09** | 1.22 |
| | | $\times 10^{-4}$ | (4.18) | (1.44) | (1.40) | (1.42) | (1.39) | (1.39) |
| | MAPE | $\times 10^{-2}$ | 626.22 | **6.67** | 6.78 | 6.77 | 7.61 | 23.27 |
| | | $\times 10^{-3}$ | (4512.74) | (2.03) | (1.83) | (3.62) | (10.46) | (146.39) |

Moreover, we selected the set of modified peptides among the validation set to evaluate the prediction performance for modified peptides. Table 3 summarizes the validation set errors of the deep learning models on modified peptides. Comparing Table 2 and 3, modified peptides were more difficult to predict and yielded larger errors than unmodified peptides for MS2 prediction; however, this phenomenon was reversed for iRT prediction. Notably, the Fusion w.o. L1 model achieved the best performance consistently for iRT prediction. This suggests that incorporating physicochemical features into the model helps improve prediction performance.

**Table 3:** Comparison of in silico spectral library generation models trained and validated on HeLa data, where prediction errors were computed only for unseen modified peptides in validation sets. The MSE, MAE, and MAPE are reported with standard errors listed below each one. The bold font indicates the smallest number among the methods.

| Model type |  |  | Prosit | Baseline | CRNN | Fusion | Fusion w.o. L1 | Fusion w.o. scale |
| --- | --- | --- | --- | --- | --- | --- | --- | --- |
| MS2<br>(+2) | MSE | $\times 10^{-4}$ | 16.90 | 6.10 | 3.45 | 4.60 | **3.39** | 3.56 |
| | | $\times 10^{-6}$ | (21.13) | (8.18) | (6.30) | (8.85) | (6.31) | (6.46) |
| | MAE | $\times 10^{-3}$ | 4.91 | 2.82 | **2.05** | 2.20 | 2.08 | 2.09 |
| | | $\times 10^{-5}$ | (3.18) | (1.91) | (1.44) | (1.66) | (1.43) | (1.46) |
| | MAPE | $\times 10^{-2}$ | 3.07 | 1.36 | 1.44 | **1.25** | 2.53 | 1.83 |
| | | $\times 10^{-5}$ | (13.25) | (8.56) | (8.62) | (7.98) | (11.73) | (9.84) |
| MS2<br>(+3) | MSE | $\times 10^{-4}$ | 24.40 | 9.58 | **5.31** | 5.70 | 6.54 | 5.87 |
| | | $\times 10^{-6}$ | (31.57) | (13.43) | (9.59) | (10.67) | (12.02) | (10.83) |
| | MAE | $\times 10^{-3}$ | 7.29 | 4.06 | **2.98** | 3.04 | 3.19 | 3.10 |
| | | $\times 10^{-5}$ | (4.99) | (3.06) | (2.28) | (2.36) | (2.53) | (2.39) |
| | MAPE | $\times 10^{-2}$ | 4.87 | **1.89** | 1.96 | 2.45 | 2.80 | 3.06 |
| | | $\times 10^{-5}$ | (22.13) | (13.05) | (12.85) | (14.57) | (15.75) | (16.47) |
| iRT | MSE | $\times 10^{-4}$ | 352.54 | 5.50 | 5.09 | 5.37 | **4.78** | 5.07 |
| | | $\times 10^{-5}$ | (57.06) | (9.35) | (8.99) | (8.87) | (8.85) | (8.70) |
| | MAE | $\times 10^{-2}$ | 18.02 | 1.11 | 1.18 | 1.19 | **1.07** | 1.19 |
| | | $\times 10^{-4}$ | (8.71) | (3.35) | (3.12) | (3.22) | (3.09) | (3.10) |
| | MAPE | $\times 10^{-2}$ | 89.19 | 4.44 | 4.67 | 4.52 | **4.31** | 4.84 |
| | | $\times 10^{-3}$ | (87.87) | (2.86) | (2.27) | (2.29) | (2.46) | (2.89) |

#### Mouse Cerebellum Data

Table 4 presents the prediction performance of deep neural network models evaluated on the Mouse cerebellum data. Similar to the results for the HeLa data, the Fusion w.o. L1 and CRNN models achieved the lowest MSE and MAE values, while the Fusion model yielded the lowest MAPE values among MS2 prediction models. This result demonstrates the overall high accuracy of the Fusion w.o. L1 and CRNN models, as well as the robustness provided by the L1-norm regularization in the Fusion model. For iRT prediction, on the other hand, the Fusion w.o. L1 and Fusion w.o. scale achieved the best performance, suggesting that the proposed fusion models using atom count features and global physicochemical features are well-suitable for iRT prediction.

**Table 4:**
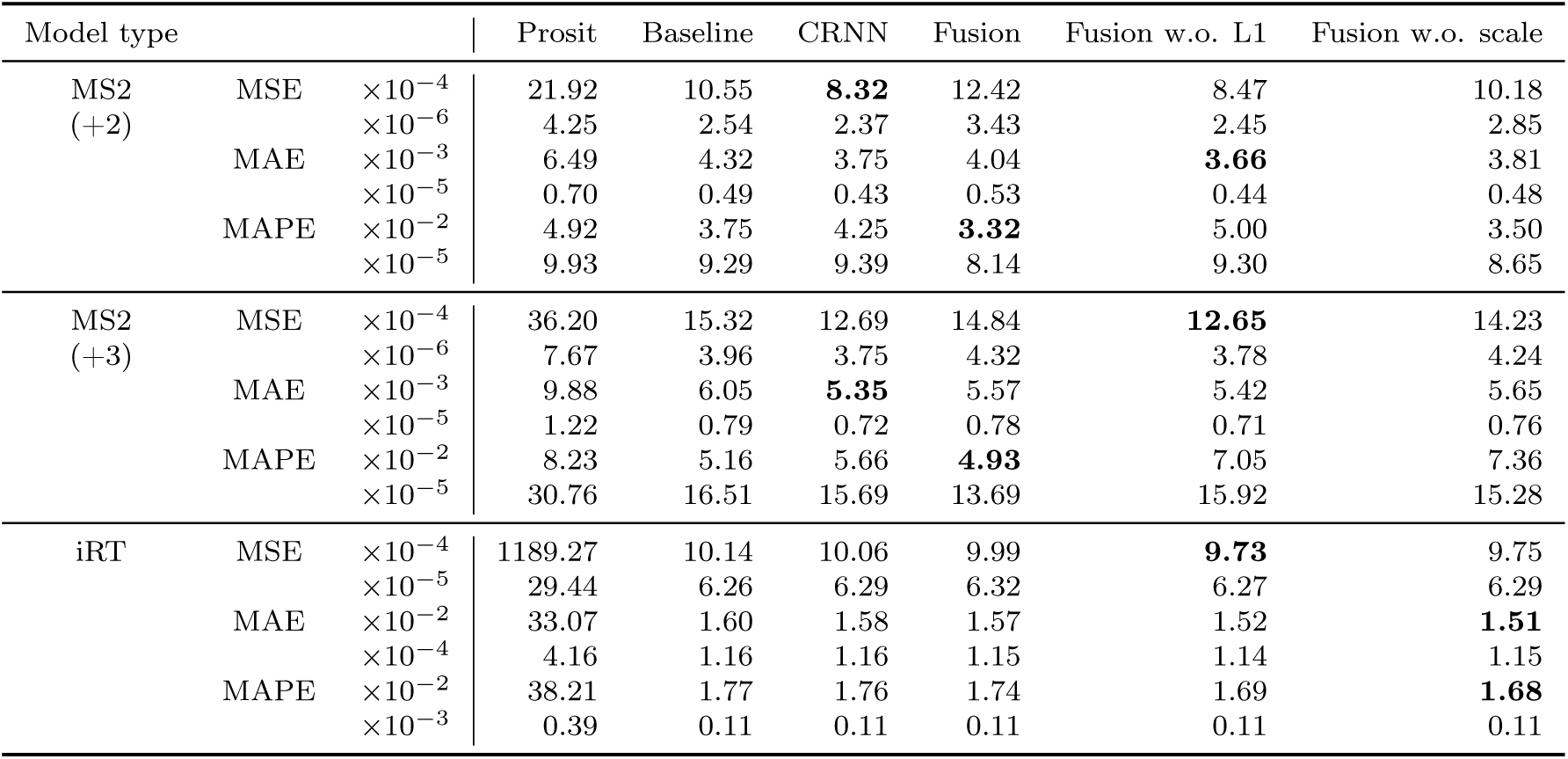
Comparison of in silico spectral library generation models trained and validated on Mouse cerebellum data. The MSE, MAE, and MAPE are reported with standard errors listed below each one. The bold font indicates the smallest number among the models.

| Model type |  |  | Prosit | Baseline | CRNN | Fusion | Fusion w.o. L1 | Fusion w.o. scale |
| --- | --- | --- | --- | --- | --- | --- | --- | --- |
| MS2<br>(+2) | MSE | $\times 10^{-4}$ | 21.92 | 10.55 | **8.32** | 12.42 | 8.47 | 10.18 |
| | | $\times 10^{-6}$ | 4.25 | 2.54 | 2.37 | 3.43 | 2.45 | 2.85 |
| | MAE | $\times 10^{-3}$ | 6.49 | 4.32 | 3.75 | 4.04 | **3.66** | 3.81 |
| | | $\times 10^{-5}$ | 0.70 | 0.49 | 0.43 | 0.53 | 0.44 | 0.48 |
| | MAPE | $\times 10^{-2}$ | 4.92 | 3.75 | 4.25 | **3.32** | 5.00 | 3.50 |
| | | $\times 10^{-5}$ | 9.93 | 9.29 | 9.39 | 8.14 | 9.30 | 8.65 |
| MS2<br>(+3) | MSE | $\times 10^{-4}$ | 36.20 | 15.32 | 12.69 | 14.84 | **12.65** | 14.23 |
| | | $\times 10^{-6}$ | 7.67 | 3.96 | 3.75 | 4.32 | 3.78 | 4.24 |
| | MAE | $\times 10^{-3}$ | 9.88 | 6.05 | **5.35** | 5.57 | 5.42 | 5.65 |
| | | $\times 10^{-5}$ | 1.22 | 0.79 | 0.72 | 0.78 | 0.71 | 0.76 |
| | MAPE | $\times 10^{-2}$ | 8.23 | 5.16 | 5.66 | **4.93** | 7.05 | 7.36 |
| | | $\times 10^{-5}$ | 30.76 | 16.51 | 15.69 | 13.69 | 15.92 | 15.28 |
| iRT | MSE | $\times 10^{-4}$ | 1189.27 | 10.14 | 10.06 | 9.99 | **9.73** | 9.75 |
| | | $\times 10^{-5}$ | 29.44 | 6.26 | 6.29 | 6.32 | 6.27 | 6.29 |
| | MAE | $\times 10^{-2}$ | 33.07 | 1.60 | 1.58 | 1.57 | 1.52 | **1.51** |
| | | $\times 10^{-4}$ | 4.16 | 1.16 | 1.16 | 1.15 | 1.14 | 1.15 |
| | MAPE | $\times 10^{-2}$ | 38.21 | 1.77 | 1.76 | 1.74 | 1.69 | **1.68** |
| | | $\times 10^{-3}$ | 0.39 | 0.11 | 0.11 | 0.11 | 0.11 | 0.11 |

#### Pombe Data Processed with Chymotrypsin

Table 5 presents the prediction performance of the deep neural network models evaluated on the Pombe (chymotrypsin) data. In the table, the Fusion w.o. L1 and CRNN models achieved the lowest MSE and MAE values in MS2(+2) prediction, which is consistent with the previous results. Similarly, in MS2(+3) prediction, the Fusion w.o. L1 model achieved the lowest MSE value and the CRNN model achieved the lowest MAE and MAPE values. The Fusion w.o. L1 model consistently achieved the highest performance in iRT prediction, demonstrating the critical role of physicochemical features in accurate iRT estimation.

**Table 5:** Comparison of in silico spectral library generation models trained and validated on Pombe (chymotrypsin) data. The MSE, MAE, and MAPE are reported with standard errors listed below each one. The bold font indicates the smallest number among the models.

| Model type |  |  | Prosit | Baseline | CRNN | Fusion | Fusion w.o. L1 | Fusion w.o. scale |
| --- | --- | --- | --- | --- | --- | --- | --- | --- |
| MS2<br>(+2) | MSE | $\times 10^{-4}$ | 61.04 | 21.81 | **18.24** | 18.51 | 18.27 | 18.42 |
| | | $\times 10^{-6}$ | 65.65 | 27.51 | 24.83 | 25.34 | 25.18 | 25.34 |
| | MAE | $\times 10^{-3}$ | 12.02 | 7.60 | 6.73 | 7.06 | **6.70** | 6.84 |
| | | $\times 10^{-5}$ | 8.67 | 5.22 | 4.78 | 4.81 | 4.79 | 4.80 |
| | MAPE | $\times 10^{-2}$ | 10.15 | 6.80 | **5.69** | 13.72 | 6.26 | 6.72 |
| | | $\times 10^{-5}$ | 267.45 | 57.40 | 42.00 | 50.31 | 42.04 | 42.76 |
| MS2<br>(+3) | MSE | $\times 10^{-4}$ | 72.06 | 28.44 | 26.14 | 25.71 | **25.51** | 25.69 |
| | | $\times 10^{-6}$ | 118.84 | 59.40 | 54.63 | 54.90 | 53.85 | 55.01 |
| | MAE | $\times 10^{-3}$ | 13.50 | 8.20 | **7.72** | 7.95 | 8.32 | 7.83 |
| | | $\times 10^{-5}$ | 15.03 | 9.46 | 9.07 | 8.99 | 8.94 | 8.99 |
| | MAPE | $\times 10^{-2}$ | 22.53 | 8.91 | **6.77** | 10.03 | 11.15 | 8.53 |
| | | $\times 10^{-5}$ | 1619.23 | 227.18 | 162.52 | 117.57 | 151.78 | 125.68 |
| iRT | MSE | $\times 10^{-4}$ | 109.03 | 10.08 | 9.14 | 9.43 | **9.07** | 9.10 |
| | | $\times 10^{-5}$ | 31.94 | 12.61 | 12.06 | 12.60 | 12.50 | 12.31 |
| | MAE | $\times 10^{-2}$ | 8.512 | 2.18 | 2.06 | 2.09 | **2.05** | 2.05 |
| | | $\times 10^{-4}$ | 14.11 | 5.52 | 5.29 | 5.39 | 5.28 | 5.30 |
| | MAPE | $\times 10^{-2}$ | 13.84 | 3.36 | 3.16 | 3.21 | **3.15** | **3.15** |
| | | $\times 10^{-3}$ | 2.84 | 1.00 | 0.97 | 0.99 | 0.97 | 0.97 |

Table 6 presents a comparison of the performance of deep neural network models on modified peptides from the Combe (chymotrypsin) dataset. Similar to Table 5, the Fusion w.o. L1 model achieved the lowest MSE and MAE values in MS2(+2) prediction, while the CRNN obtained the lowest MAE and MAPE values in MS2(+3) prediction. Additionally, the Fusion w.o. L1 model outperformed others by yielding the lowest MSE, MAE, and MAPE values in iRT prediction. Notably, the Baseline model achieved the lowest MAPE value in MS2(+2) prediction and the lowest MSE value in MS2(+3) prediction. This suggests that its simple model architecture may have reduced overfitting in small datasets, while the incorporation of physicochemical features in the Fusion w.o. L1 model helped prevent overfitting and improved generalization.

**Table 6:** Comparison of in silico spectral library generation models trained and validated on Pombe (chymotrypsin) data, where prediction errors were computed only for unseen modified peptides in validation sets. The MSE, MAE, and MAPE are reported with standard errors listed below each one. The bold font indicates the smallest number among the methods.

| Model type |  |  | Prosit | Baseline | CRNN | Fusion | Fusion w.o. L1 | Fusion w.o. scale |
| --- | --- | --- | --- | --- | --- | --- | --- | --- |
| MS2<br>(+2) | MSE | $\times 10^{-4}$ | 73.28 | 31.55 | 31.25 | 31.21 | **31.07** | 31.41 |
| | | $\times 10^{-6}$ | 563.36 | 396.82 | 394.85 | 395.92 | 399.58 | 400.96 |
| | MAE | $\times 10^{-3}$ | 13.68 | **9.05** | 9.11 | 9.30 | **9.05** | 9.18 |
| | | $\times 10^{-5}$ | 72.66 | 63.40 | 63.08 | 63.01 | 62.90 | 63.23 |
| | MAPE | $\times 10^{-2}$ | 11.54 | **7.06** | 7.99 | 15.44 | 8.71 | 9.04 |
| | | $\times 10^{-5}$ | 1498.91 | 432.82 | 749.35 | 665.44 | 692.01 | 675.15 |
| MS2<br>(+3) | MSE | $\times 10^{-4}$ | 78.73 | **40.49** | 41.80 | 42.85 | 41.43 | 42.06 |
| | | $\times 10^{-6}$ | 761.07 | 644.46 | 664.96 | 680.45 | 655.31 | 659.29 |
| | MAE | $\times 10^{-3}$ | 15.16 | 10.41 | **9.84** | 10.49 | 10.84 | 10.40 |
| | | $\times 10^{-5}$ | 99.99 | 97.84 | 99.60 | 100.72 | 98.89 | 99.78 |
| | MAPE | $\times 10^{-2}$ | 10.90 | 6.04 | **5.13** | 9.35 | 10.08 | 7.96 |
| | | $\times 10^{-5}$ | 1126.31 | 426.00 | 361.85 | 463.53 | 520.56 | 433.48 |
| iRT | MSE | $\times 10^{-4}$ | 134.64 | 55.52 | 48.04 | 56.97 | **47.92** | 52.19 |
| | | $\times 10^{-5}$ | 179.63 | 218.20 | 188.65 | 204.60 | 190.48 | 192.05 |
| | MAE | $\times 10^{-2}$ | 9.29 | 5.28 | 4.93 | 5.50 | **4.83** | 5.20 |
| | | $\times 10^{-4}$ | 62.17 | 69.08 | 63.99 | 67.84 | 65.14 | 65.79 |
| | MAPE | $\times 10^{-2}$ | 14.14 | 8.61 | 8.11 | 8.93 | **7.95** | 8.53 |
| | | $\times 10^{-3}$ | 11.49 | 14.98 | 14.26 | 14.57 | 14.45 | 14.32 |

#### Pombe Data Processed with Trypsin and Chymotrypsin

Tables 7 and 8 present the performance of deep neural network models on the Pombe (trypsin-chymotrypsin) dataset. This dataset is slightly larger than the Pombe (chymotrypsin) dataset, as sequential digestion with two proteases improved peptide and protein identification^36^. While the Fusion w.o. L1 model achieved the lowest MSE, MAE, and MAPE in iRT prediction, it is notable that the Fusion w.o. scale model showed the second-best performance. This suggests that physicochemical features other than amino acid scale features– such as atom count and global features–also contributed to improved iRT prediction.

**Table 7:** Comparison of in silico spectral library generation models trained and validated on Pombe (trypsin-chymotrypsin) data. The MSE, MAE, and MAPE are reported with standard errors listed below each one. The bold font indicates the smallest number among the models.

| Model type |  |  | Prosit | Baseline | CRNN | Fusion | Fusion w.o. L1 | Fusion w.o. scale |
| --- | --- | --- | --- | --- | --- | --- | --- | --- |
| MS2<br>(+2) | MSE | $\times 10^{-4}$ | 51.03 | 15.65 | **10.61** | 11.93 | 10.88 | 11.24 |
| | | $\times 10^{-6}$ | 31.60 | 12.03 | 9.49 | 10.59 | 9.93 | 10.07 |
| | MAE | $\times 10^{-3}$ | 10.55 | 6.01 | 4.95 | 4.95 | **4.81** | 5.04 |
| | | $\times 10^{-5}$ | 4.28 | 2.41 | 1.98 | 2.10 | 2.01 | 2.04 |
| | MAPE | $\times 10^{-2}$ | 11.18 | 6.22 | 5.81 | **4.71** | 5.21 | 5.91 |
| | | $\times 10^{-5}$ | 192.04 | 36.02 | 27.58 | 25.80 | 27.02 | 31.15 |
| MS2<br>(+3) | MSE | $\times 10^{-4}$ | 89.38 | 26.83 | 23.27 | **22.93** | 23.00 | 23.60 |
| | | $\times 10^{-6}$ | 77.69 | 31.61 | 28.91 | 28.58 | 28.65 | 29.23 |
| | MAE | $\times 10^{-3}$ | 16.43 | 8.56 | **7.73** | 8.19 | 7.86 | 7.93 |
| | | $\times 10^{-5}$ | 9.92 | 5.49 | 5.12 | 5.07 | 5.09 | 5.15 |
| | MAPE | $\times 10^{-2}$ | 39.08 | 8.61 | **7.93** | 12.48 | 9.16 | 9.84 |
| | | $\times 10^{-5}$ | 2290.34 | 66.70 | 61.12 | 66.38 | 59.48 | 87.69 |
| iRT | MSE | $\times 10^{-4}$ | 134.14 | 5.70 | 5.77 | 5.63 | **4.66** | 4.80 |
| | | $\times 10^{-5}$ | 23.63 | 4.94 | 5.41 | 5.05 | 4.89 | 4.86 |
| | MAE | $\times 10^{-2}$ | 9.47 | 1.44 | 1.41 | 1.42 | **1.26** | 1.29 |
| | | $\times 10^{-4}$ | 8.61 | 2.55 | 2.61 | 2.55 | 2.35 | 2.38 |
| | MAPE | $\times 10^{-2}$ | 16.21 | 2.26 | 2.21 | 2.23 | **1.97** | 2.02 |
| | | $\times 10^{-3}$ | 1.83 | 0.39 | 0.40 | 0.40 | 0.36 | 0.37 |

**Table 8:** Comparison of in silico spectral library generation models trained and validated on Pombe (trypsin-chymotrypsin) data, where prediction errors were computed only for unseen modified peptides in validation sets. The MSE, MAE, and MAPE are reported with standard errors listed below each one. The bold font indicates the smallest number among the methods.

| Model type |  |  | Prosit | Baseline | CRNN | Fusion | Fusion w.o. L1 | Fusion w.o. scale |
| --- | --- | --- | --- | --- | --- | --- | --- | --- |
| MS2<br>(+2) | MSE | $\times 10^{-4}$ | 57.78 | 23.38 | **19.20** | 20.57 | 19.70 | 20.17 |
| | | $\times 10^{-6}$ | 285.58 | 191.64 | 173.80 | 184.67 | 179.72 | 181.90 |
| | MAE | $\times 10^{-3}$ | 11.75 | 7.24 | 6.50 | 6.44 | **6.37** | 6.64 |
| | | $\times 10^{-5}$ | 37.84 | 31.17 | 28.26 | 29.28 | 28.64 | 28.96 |
| | MAPE | $\times 10^{-2}$ | 12.70 | 7.15 | 7.24 | **5.92** | 6.53 | 7.58 |
| | | $\times 10^{-5}$ | 1114.56 | 496.99 | 317.39 | 286.09 | 299.89 | 353.35 |
| MS2<br>(+3) | MSE | $\times 10^{-4}$ | 101.92 | 36.18 | **34.45** | 34.72 | 34.68 | 34.85 |
| | | $\times 10^{-6}$ | 426.45 | 239.01 | 239.25 | 239.32 | 237.64 | 240.47 |
| | MAE | $\times 10^{-3}$ | 19.64 | 11.52 | **11.10** | 11.66 | 11.25 | 11.13 |
| | | $\times 10^{-5}$ | 55.57 | 41.76 | 40.76 | 40.85 | 40.88 | 41.00 |
| | MAPE | $\times 10^{-2}$ | 27.10 | 9.39 | **9.35** | 13.86 | 10.00 | 10.21 |
| | | $\times 10^{-5}$ | 4055.51 | 368.63 | 262.28 | 285.73 | 256.81 | 260.89 |
| iRT | MSE | $\times 10^{-4}$ | 151.01 | 26.41 | 28.02 | 27.29 | **9.26** | 12.99 |
| | | $\times 10^{-5}$ | 79.17 | 21.15 | 22.12 | 22.91 | 19.38 | 20.41 |
| | MAE | $\times 10^{-2}$ | 10.13 | 4.53 | 4.68 | 4.59 | **2.16** | 2.76 |
| | | $\times 10^{-4}$ | 30.64 | 15.75 | 16.02 | 16.13 | 13.91 | 15.02 |
| | MAPE | $\times 10^{-2}$ | 16.21 | 7.35 | 7.69 | 7.43 | **3.42** | 4.48 |
| | | $\times 10^{-3}$ | 5.45 | 2.79 | 2.91 | 2.81 | 2.30 | 2.59 |

### Comparison of Spectral Libraries Generated by Deep Learning Models

This section compares the spectral libraries generated by the deep learning models. An in silico spectral library can be generated by combining an MS2 prediction model and an iRT prediction model. We combined the same types of the MS2 and iRT prediction models as listed in Section Hyperparameter Optimization.

In this study, we prepared two separate lists of peptides to predict MS2 and iRT values. One is the peptide list from the Pan-Human spectral library^37^, and the other is the union of the peptide lists from the Pan-Human spectral library and HeLa cell-based spectral library, which has been mentioned in previous paragraphs. We can see that the in silico spectral libraries can predict spectra of the peptides that have not been included in the HeLa-cell based experimental library.Table 9 compares the spectral libraries generated by deep learning models. Each row of the table represents quantitative metrics for the comparison. The spectral libraries were compared in terms of the number of unique peptide precursors including charge (Precursors), the number of unique stripped sequences (Peptides), the number of unique stripped sequences which point to only one protein (Proteotypic Peptides), the number of unique protein groups identified by a single peptide precursor (Single Hits), the number of fragment ions (Fragments), and the total number of fragment ions (Fragments Total). In Table 9, we can find that the Fusion w.o. L1 models yielded the largest numbers of peptides and fragments, and the Fusion model yielded the least numbers. This result implies that neural network weights suppressed by the L1-norm regularization affected the reduced number of fragment ions.

**Table 9:** Comparison of spectral libraries generated by deep learning models regarding the number of peptides, proteins, and fragments included. The HeLa-cell based experimental spectral library used as the training dataset is listed in the right-most column for comparison. The spectral libraries generated by the deep learning models were imported and filtered using Spectronaut based on default settings. The quantitative metrics in the table rows are unique to MS2 prediction models.

| Pan-Human | Baseline | CRNN | Fusion | Fusion w.o. L1 | Fusion w.o. scale | Experimental |
| --- | --- | --- | --- | --- | --- | --- |
| Precursors | 167,046 | 167,139 | 167,133 | 167,142 | 167,141 | 96,354 |
| Peptides | 135,911 | 135,976 | 135,976 | 135,979 | 135,976 | 69,515 |
| Proteotypic Peptides | 129,707 | 129,769 | 129,770 | 129,772 | 129,769 | 65,888 |
| Protein Groups | 10,308 | 10,308 | 10,308 | 10,308 | 10,308 | 6,088 |
| Proteins | 10,452 | 10,452 | 10,452 | 10,452 | 10,452 | 6,180 |
| Single Hits | 1,274 | 1,273 | 1,273 | 1,273 | 1,273 | 546 |
| Fragments | 991,550 | 996,586 | 989,142 | 997,912 | 995,854 | 567,434 |
| Fragments Total | 2,110,432 | 2,618,537 | 2,397,971 | 2,542,664 | 2,597,544 | 1,187,745 |
| Pan-Human + HeLa | Baseline | CRNN | Fusion | Fusion w.o. L1 | Fusion w.o. scale | Experimental |
| Precursors | 190,813 | 190,911 | 190,895 | 190,920 | 190,915 | 96,354 |
| Peptides | 151,131 | 151,199 | 151,190 | 151,204 | 151,199 | 69,515 |
| Proteotypic Peptides | 144,195 | 144,257 | 144,250 | 144,262 | 144,257 | 65,888 |
| Protein Groups | 10,444. | 10,444 | 10,444 | 10,444 | 10,444 | 6,088 |
| Proteins | 10,586 | 10,586 | 10,586 | 10,586 | 10,586 | 6,180 |
| Single Hits | 1,215 | 1,214 | 1,214 | 1,214 | 1,214 | 546 |
| Fragments | 1,133,818 | 1,138,566 | 1,130,840 | 1,139,519 | 1,137,566 | 567,434 |
| Fragments Total | 3,088,284 | 3,809,804 | 3,455,875 | 3,720,513 | 3,775,313 | 1,187,745 |

### DIA Data Analysis Performance Comparison

In this section, we compare in silico-generated spectral libraries in terms of peptide and protein identification using benchmark DIA datasets. Raw DIA datasets and generated in silico spectral libraries were imported and analyzed using Spectronaut 16 with default settings. Based on deep learning models trained on the DDA data of HeLa cells, two types of spectral libraries were generated: one is for the peptides from the Pan-Human spectral library and the other for peptides from the Pan-Human and HeLa spectral libraries.

The upper panel of Table 10 summarizes the numbers of identified peptides and proteins when the spectral libraries were generated for the peptides contained in the Pan-Human spectral library. Among the deep learning methods compared, the CRNN model yielded the greatest numbers of identified peptides, and the Fusion w.o. scale model yielded the greatest numbers of proteins. Notably, the Fusion model yielded the least numbers of identified peptides and proteins, implying that regularizing neural networks for amino acid scale features might have adversely affected the identification of peptides and proteins. And the greatest numbers of identified proteins yielded by the Fusion w.o. scale model demonstrates that the other physicochemical features processed by regularized neural networks were helpful for improved identification of proteins.

**Table 10:** Comparison of spectral libraries regarding the numbers of identified peptides and proteins from HeLa cell DIA data. In this experiment, deep learning models were used to generate in silico spectral libraries for the peptides contained either in the Pan-Human spectral library or in the Pan-Human and HeLa cell-based experimental spectral library. The bold font indicates the largest number among different models.

| Pan-Human | Baseline | CRNN | Fusion | Fusion w.o. L1 | Fusion w.o. scale |
| --- | --- | --- | --- | --- | --- |
| Precursors | 69,875 | **70,695** | 69,332 | 70,470 | 69,540 |
| Peptides | 58,855 | **59,579** | 58,502 | 59,345 | 58,648 |
| Protein Groups | 5,953 | 5,932 | 5,892 | 5,994 | **6,050** |
| Proteins | 6,014 | 5,996 | 5,952 | 6,060 | **6,121** |
| Pan-Human + HeLa | Baseline | CRNN | Fusion | Fusion w.o. L1 | Fusion w.o. scale |
| Precursors | 85,542 | 86,261 | 85,156 | **87,187** | 85,479 |
| Peptides | 70,259 | 70,798 | 70,040 | **71,610** | 70,288 |
| Protein Groups | 6,142 | 6,155 | 6,112 | **6,303** | 6,150 |
| Proteins | 6,201 | 6,212 | 6,165 | **6,367** | 6,207 |

The lower panel of Table 10 presents the numbers of identified peptides and proteins obtained by in silico spectral libraries generated for peptides contained in the union of the Pan-Human and HeLa experimental spectral libraries. Comparing the upper and lower panels, we can find that including the peptides in the HeLa experimental spectral library increased the numbers of identified peptides and proteins significantly. Comparing the deep learning models, we can find that the Fusion w.o. L1 model yielded the largest numbers of identified peptides and proteins, while the Fusion model yielded the smallest numbers. This implies that while the L1-norm regularization on physicochemical features might have been helpful in the identification of proteins in the Pan-Human spectral library, it adversely affected in the HeLa experimental spectral library.

Figure 3 illustrates the numbers of identified peptides and protein groups by the in silico spectral libraries generated by the Baseline, CRNN, and Fusion w.o. scale models, where the peptide list of the libraries was retrieved from the Pan-Human spectral library. The results from the other models were omitted for simplicity. The lower panels of the figure show that a large number of peptides are shared by the CRNN and Fusion w.o. scale models and a large number of proteins are shared by the Baseline and Fusion w.o. scale model. Moreover, it is shown that the Fusion w.o. scale model identified the largest number of protein groups, demonstrating the competitiveness of the proposed model.

**Figure 3.**
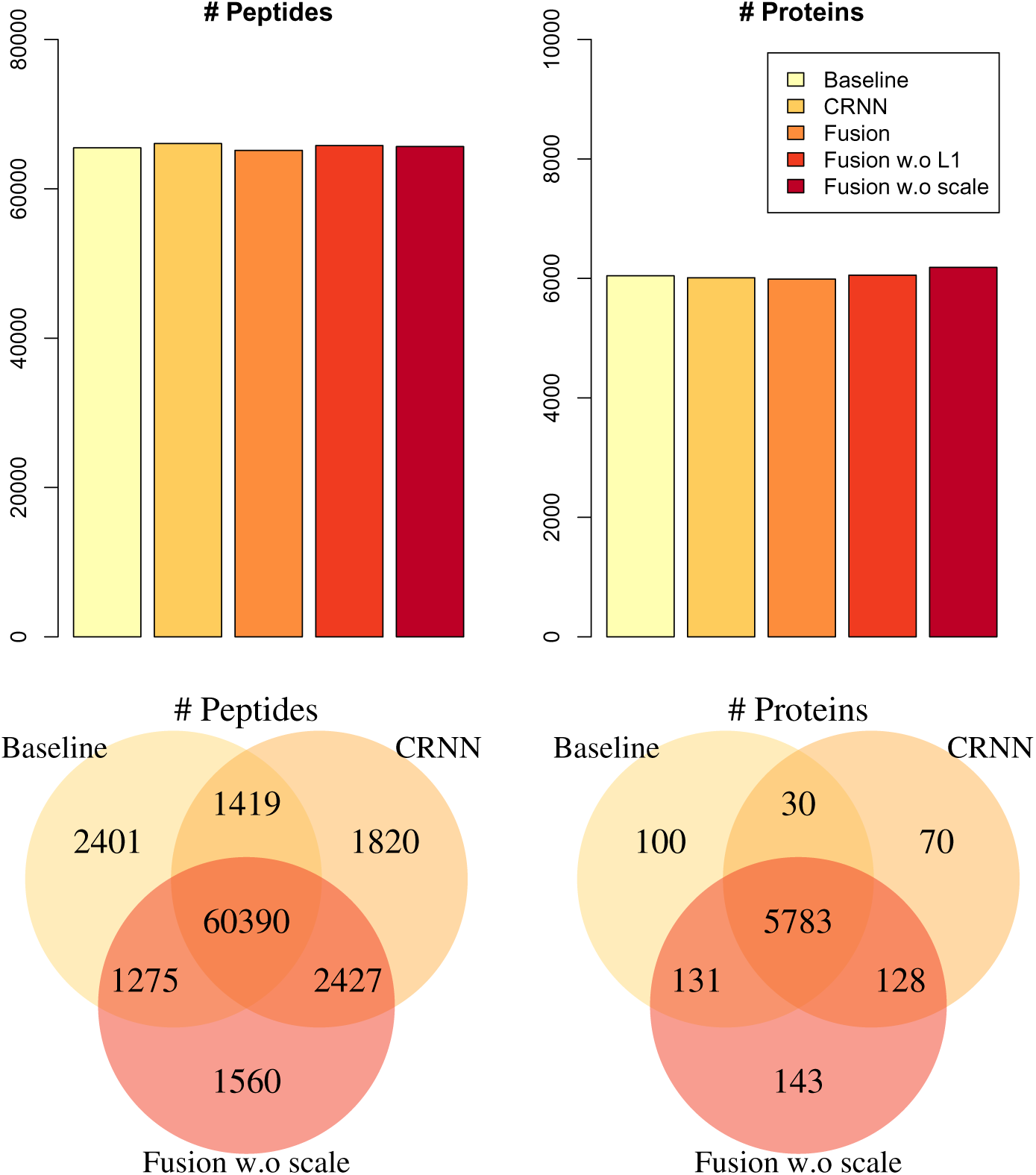
: Number of identified peptides (left panels) and protein groups (right panels) from HeLa cell DIA data, where in silico spectral libraries were generated for the peptides contained in the Pan-Human spectral library.

Figure 4 illustrates the numbers of identified peptides and protein groups by the in silico spectral libraries generated by the Baseline, CRNN, and Fusion w.o. L1 models, where the peptide list of the libraries was retrieved from the Pan-Human and HeLa experimental libraries. The Fusion w.o. L1 model was selected because it showed the largest numbers of identified peptides and proteins in the lower panel of Table 10. The lower panels of the figure show that more peptides and protein groups are shared by the CRNN and Fusion w.o. L1 models, and the Fusion w.o. L1 model identified the largest number of protein groups.

**Figure 4.**
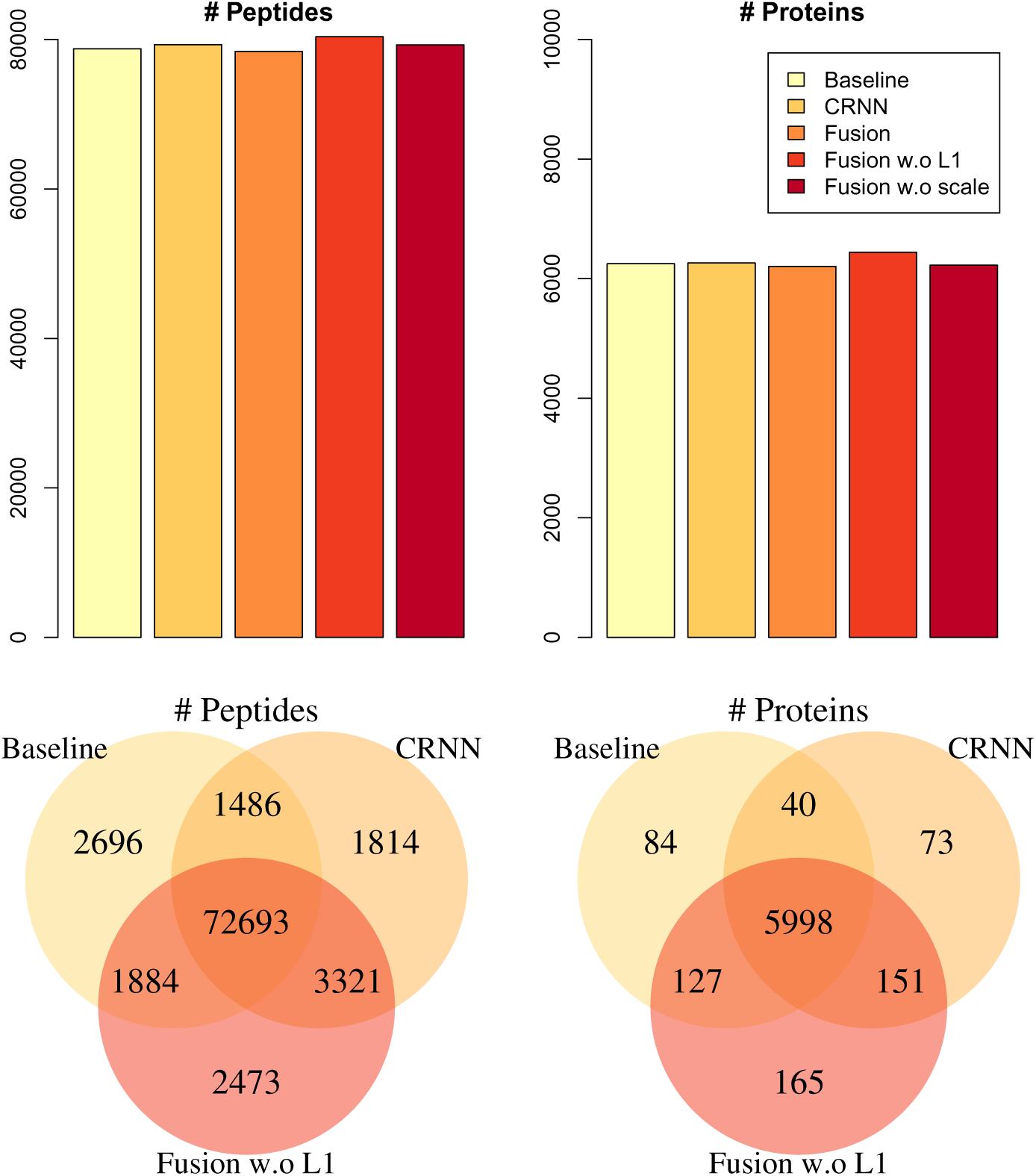
: Number of identified peptides (left panels) and proteins (right panels) from HeLa cell DIA data, where in silico spectral libraries were generated for the peptides contained in the Pan-Human and the HeLa experimental spectral libraries.

Notably, relatively large numbers of peptides were identified uniquely by the Baseline model in Figures 3 and 4, however, the Fusion w.o. scale and Fusion w.o. L1 models identified larger numbers of proteins uniquely. This implies that the Fusion models could identify more peptides which were important for identifying proteins than the Baseline model.

## Discussion and Conclusions

This study proposes a deep learning-based method for in silico spectral library generation. The proposed method effectively combines traditional one-hot encoding features with the physicochemical features of peptides by employing L1-norm regularization and a transfer learning strategy. The physicochemical features used in this study are categorized into vector-valued and matrix-valued types. Vector-valued features represent the global features of the peptide as a whole, whereas matrix-valued features capture characteristics extracted based on the sequence of amino acids constituting the peptide. The regularized neural networks used in this study are designed to process both types of features in parallel and integrate them through fully connected layers. Furthermore, the proposed transfer learning strategy improves the training efficiency by dividing a high-dimensional optimization problem into two separate, lower dimensional problems.

In numerical experiments using data obtained from cell lines of different species, including human, mouse, and S. pombe, the proposed method outperformed existing deep learning methods such as Prosit and DeepDIA. Specifically, the results showed that hyperparameter optimization led to well-tuned deep neural network architectures, resulting in superior performance by the CRNN and the family of Fusion models. Furthermore, comparison with model variants–featuring slightly different combinations of input features and hyperparameters–demonstrated that the Fusion w.o. L1 and CRNN models achieved the lowest MSE and MAE values in most MS2 prediction tasks, and the Fusion w.o. L1 model consistently showed the lowest MSE, MAE, and MAPE values in most iRT prediction tasks. These results imply that physicochemical features provide a significant advantage for iRT prediction and that incorporating them into the model architecture does not degrade MS2 prediction performance.

The DIA data analysis showed that the Fusion w.o. scale model identified the highest number of proteins using peptides from Pan-Human library. This result suggests that atom count and global features may aid in protein group identification, particularly when regularized with an L1-norm. In contrast, the Fusion model identified the fewest peptides and proteins, while the Fusion w.o. L1 model yielded identified the most peptides and proteins using peptides from the combined Pan-Human + HeLa library. These findings indicate that incorporating physicochemical features is beneficial for DIA data analysis. However, applying L1-norm regularization, especially to amino acid scale features, may negatively affect performance.

While the proposed method incorporates various physicochemical features of peptides, many other factors that may influence tandem MS measurement were not considered in this study. For example, difference in MS instruments and experimental settings, such as collision energy and collision cell type, can lead to variations in MS2 spectra ^15^. By incorporating features related to these experimental conditions, the proposed deep learning model could potentially improve its predictive performance across diverse datasets generated under varying conditions. In future work, we plan tov extend the proposed method to account for different instrument types, experimental settings, peptide modifications, and diverse cell lines and species.

## Acknowledgement

This work was supported by National Research Foundation of Korea (NRF) grants funded by the Korea government (MSIT) (Nos. RS-2021-NR061647, RS-2024-00358572, RS-2024-00336424).

## Supporting Information Available

- supporting-information.pdf: deep learning model architectures

